# Spatiotemporal patterns of dunlin (*Calidris alpina*) in continental lakes of the Iberian Peninsula

**DOI:** 10.1101/434886

**Authors:** M. S. S. Gonçalves, J. A. Gil-Delgado, R. U. Gosálvez, G. M. López-Iborra, A. Ponz, P. S. Pons, A. Velasco

## Abstract

Spatiotemporal dynamics may present different levels of regional or local stability, generally attributed to local habitat and landscape factors, reflecting the tolerances and ecological requirements of the populations. In this study, we examined the variations of dunlin abundance and occurrence in twenty-three wetlands of the “La Mancha Húmeda” Biosphere Reserve, central Spain, between October 2010 and July 2017. In addition, we observed the variations of local abundance in the lakes of the Manjavacas lagoon complex, seeking to understand the factors that determine the local movements of the wintering individuals. Eleven lakes had records of dunlin, but most of individuals (ca. 90%) were observed in the Manjavacas Lagoon Complex (Alcahozo and Manjavacas lakes). The preference for this complex associated with temporal variations of abundance between the Alcahozo and Manjavacas lagoons possibly reflects the specific characteristics of the invertebrate community available as prey, especially for the presence of anostracans and copepods. The results of this study are a first step in understanding the factors that lead this species to select continental wetlands as wintering sites.

## 1. Introduction

Spatiotemporal dynamics may present different levels of regional or local stability, generally attributed to habitat and landscape factors, reflecting the tolerances and ecological requirements of the populations (Collins & Glenn, 1991). In waterbirds, the spatiotemporal distribution in wetland patches has often been associated to destruction and environmental fragmentation (Guadagnin & Maltchik, 2007). Inland wetland ecosystems are distributed in spots on landscape. Therefore, in order to maintain the stability of the populations, waterbirds are forced to adapt to local conditions of habitat change as well as incorporating multiple fragments or habitat patches into the daily life cycle (Guadagnin et al. 2009). Among waterbird migratory species, the recognition of these variations helps to understand the relevance of interconnected habitats between breeding, wintering and stopover areas (Andres et al. 2012).

Migratory waterbirds are often used as indicators of environmental quality (Green & Elmberg, 2013). However, for many inland wetlands of high environmental importance, there is no information on spatiotemporal dynamics of migratory waders and their correlations with environmental factors (Morrison et al. 2006). This is particularly important as significant reductions in populations of many wader species have been observed over the past 30 years (Andres et al. 2012). In this context, continental wetlands require special attention since it has demonstrated a greater loss of habitat in relation to coastal environments (Kingsford et al. 2016).

In central Spain, “La Mancha Húmeda” Biosphere Reserve (hereafter MHBR) presents a complex and extensive network of permanent and temporary salt lakes of great relevance in the cycle of migratory waterbirds (Gosálvez et al. 2012). The MHBR is key habitat for those species that arrive from northern Europe towards the south of the continent European and African during the autumn and winter (BirdLife International, 2018). More than 25 wading species – approximately 40% of species in Spain, occupy the inland wetlands of the reserve (Gonçalves et al. 2018). According to the maps available at BirdLife International (2018), dunlin (*Calidris alpina*), together with black-tailed godwit (*Limosa limosa*) and common greenshank (*Tringa nebularia*), represent one of the most conspicuous winter migratory species in central Spain, however, little is known about their spatiotemporal patterns, especially in inland wetlands (Gosálvez et al. 2012; Gonçalves et al. 2016, 2018).

Determining routes of migratory birds, as well as their feeding and wintering areas, is crucial to define and implement management actions at multiple spatial scales (Andres et al. 2012). In this study, we aim to describe the variations of Dunlin abundance and occurrence in 23 wetlands of the RBMH. In addition, we observed the variations of local abundance in the lakes of the Manjavacas lagoon complex, seeking to understand the factors that determine the local movements of the wintering individuals.

## 2. Methods

### Study area

“La Mancha Húmeda” Biosphere Reserve, was created in 1981 initially with a surface of 25,000 ha (Crespo et al. 2011). Additionally, in agreement with the document of the Ministry of Agriculture, Food, and Environment (2014), this area was enlarged to about 420,000ha, inserting a large number of legally protected wetlands inside reserve. Although the relief of central Spain is marked by a great geological and geomorphological diversity, much of the reserve landscape is dominated by slightly undulating fields and extensive cereal and vineyards plantations the most important economic activity of the region (Ruiz-Pulpón, 2013; 2015). The rainfall of the region is low, varying between 300 and 500 mm annually and, combined with a warmest summer. The higher temperatures and lower rainfalls since the endo of spring, forces the lakes to a natural temporary variation in water levels (Martinez-Santos et al., 2008). Thus, from the end of July to the first rainfalls of autumn the lakes are dry. In its domains, two important conservation sites are inserted: Tablas de Daimiel National Park (approximately 2,000 hectares) and Ruidera Natural Park (3,772 hectares). Further, other wetlands are also classified as Ramsar sites and/or inserted into Natura 2000 network (Gonçalves et al. 2018).

### Bird protocol and statistical analyzes

Dunlins were counted monthly in 23 lakes between October 2010 and July 2017 (Table 1). Except for Redondilla lake, the selected lakes are concentrated in the north of the “La Mancha Húmeda” Biosphere Reserve. The 23 lakes represent in large part the high heterogeneity of the wetlands of the region, especially in relation to hydrological dynamics and in physical characteristics (Gonçalves et al. 2016). Counts were made at the end of each month between 08:00-12:00, always on favorable weather conditions (little wind or no rain). Telescopes and binoculars were used during countings.

**Table 1.** Twenty-three lakes monitored between October 2010 and July 2017 in “La Mancha Húmeda” Biosphere Reserve (see Figure 1). UTM – Datum ETRS89; Area (ha); Flooded surface (ha); Veg – area covered by natural vegetation (ha); Ni – number of sedimentary islands; IsA – total area of existing sedimentary islands (ha); AD – average depth (meters); Hyd – hydroperiod (% of months with water); Leng – shoreline length (meters) and; SDI – Shoreline Development Index.

| Code | Lake | UTM X | UTM Y | Area | Flooded | Veg | Ni | IsA | AD | Hyd | Leng | SDI |
| --- | --- | --- | --- | --- | --- | --- | --- | --- | --- | --- | --- | --- |
| 1 | Alcahozo* | 510620 | 4360164 | 88.6 | 69 | 19.5 | 0 | 0 | 1.75 | 58.5 | 4805 | 1,63 |
| 2 | Alttillo Grande | 474187 | 4393670 | 32.9 | 20.4 | 12.4 | 0 | 0 | 4.25 | 43.9 | 2932 | 1,82 |
| 3 | Artevi | 472611 | 4385210 | 27.9 | 16.7 | 11.2 | 0 | 0 | 4.75 | 68.2 | 3102 | 2,14 |
| 4 | Camino de Villafranca | 478055 | 4362616 | 159.9 | 134.7 | 23.7 | 2 | 1.7 | 2.25 | 95.1 | 9181 | 2,23 |
| 5 | Campo de Mula | 464695 | 4381452 | 50.9 | 25.5 | 25.3 | 0 | 0 | 3.25 | 21.9 | 4038 | 2,25 |
| 6 | Longar | 472448 | 4395066 | 259.5 | 99.1 | 160.3 | 0 | 0 | 4 | 92.6 | 5895 | 1,67 |
| 7 | Manjavacas | 511840 | 4363360 | 262.8 | 197.8 | 64.0 | 3 | 9 | 2.75 | 97.5 | 9872 | 1,98 |
| 8 | Mermejuela | 488191 | 4376617 | 9.9 | 8.6 | 1.2 | 0 | 0 | 6.5 | 82.9 | 1374 | 1,31 |
| 9 | Miguel Esteban | 495294 | 4373855 | 92.2 | 23.4 | 68.8 | 0 | 0 | 2.5 | 97.5 | 3966 | 2,31 |
| 10 | Pedro Muñoz | 504589 | 4362479 | 41.3 | 26.1 | 15.2 | 0 | 0 | 3.25 | 87.8 | 5936 | 3,27 |
| 11 | Peña Hueca | 470426 | 4373729 | 158.8 | 132.6 | 19.6 | 4 | 6.6 | 3.75 | 58.5 | 11146 | 2,73 |
| 12 | Quero | 478215 | 4372367 | 95.7 | 84.0 | 11.6 | 0 | 0 | 2 | 70.7 | 4991 | 1,53 |
| 13 | Retamar | 502427 | 4364111 | 79.1 | 62.7 | 16.4 | 1 | 4 | 3.5 | 21.9 | 5217 | 2,21 |
| 14 | Tírez | 469331 | 4376605 | 131.8 | 81.1 | 50.6 | 1 | 0.04 | 4.25 | 51.2 | 6112 | 1,91 |
| 15 | Veguilla | 479389 | 4360715 | 89.0 | 44.3 | 44.7 | 0 | 0 | 4.25 | 97.5 | 9607 | 4,06 |
| 16 | Larga de Villacañas | 472815 | 4384080 | 167.3 | 114.5 | 52.7 | 0 | 0 | 3.25 | 100 | 9648 | 2,54 |
| 17 | Albardiosa | 474994 | 4390229 | 71.8 | 39.5 | 32.3 | 0 | 0 | 2.5 | 8.3 | 3557 | 1,59 |
| 18 | Altillo Chica | 473975 | 4394727 | 24.6 | 17.8 | 6.8 | 0 | 0 | 3 | 55.5 | 2200 | 1,47 |
| 19 | Camino de Turleque | 464403 | 4384224 | 48.6 | 39.4 | 9.2 | 0 | 0 | 5.25 | 5.5 | 3097 | 1,39 |
| 20 | Pajares | 482321 | 4367206 | 23.5 | 20.9 | 2.6 | 0 | 0 | 2.75 | 39 | 2360 | 1,45 |
| 21 | Redondilla | 513136 | 4310021 | 5.7 | 3.5 | 2.2 | 0 | 0 | 11 | 26.8 | 981 | 1,47 |
| 22 | Salicor | 485033 | 4368502 | 81.4 | 48.3 | 23.7 | 3 | 9.2 | 5.75 | 55.5 | 5159 | 2,09 |
| 23 | Yeguas | 475657 | 4363078 | 96.2 | 64.9 | 31.3 | 0 | 0 | 2 | 61.1 | 4593 | 1,6 |

**Fig. 1.**
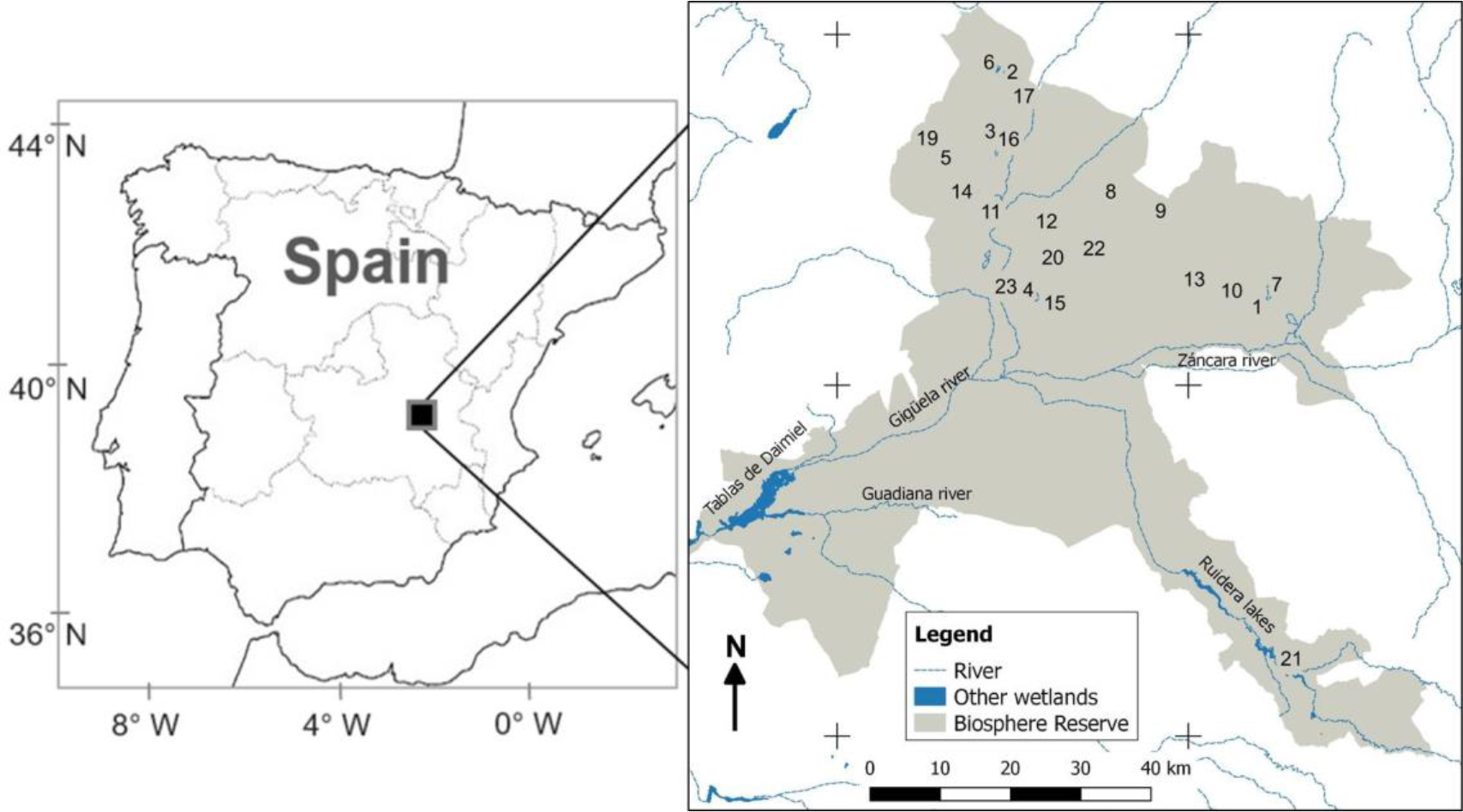
La Mancha Húmeda” Biosphere Reserve, and location of the twenty-three lakes (identification codes are in Table 1) monitored between 2010 and 2017.

Frequency of occurrence (expressed in% of months with species) and dominance (expressed in% of records in relation to total recorded individuals) were calculated for each lake in two periods: wintering (November to March) and non-wintering (April to October). The higher number of the records of individuals were concentrated in two lakes of the Manjavacas Lagoon Complex. Thus, to determine possible patterns of local movement of individuals within this area, we evaluated the level of correlation of the monthly abundance variations between the two lakes using the Pearson’s Correlation Coefficient (r). The analysis was calculated from the time series of smoothed monthly abundance of each lake. This analysis was performed using PAST software (Hammer et al. 2001). Smoothing was performed to reduce the effect of daily and monthly variability and focussing on the temporal behavior of abundance curves. Thus, the monthly data series was smoothed out by calculating the central moving average for each month using a three-month window, i.e. mean between the month, the previous month, and the subsequent month. Thus, the first and last month were excluded from the analysis.

## 3. Results

Of the 23 lakes monitored monthly between October 2010 and July 2017, eleven had records of Dunlin. A total of 5707 records were accumulated throughout the study period, with a high dominance of the individuals recorded from Alcahozo and Manjavacas lakes – approximately 90% of dominance (Table 2). During the wintering period, the Alcahozo lake had the highest average monthly abundance of individuals (81.1 ± 99.7) and the highest frequency of occurrence (54.2%). Out of the wintering period, this lake presented low values of average abundance (0.4 ± 3.1) and frequency of occurrence (4.2%). For this period, the Manjavacas and Camino de Villafranca lagoons were the most used by the species (Table 2; Fig. 1).

**Table 2.**
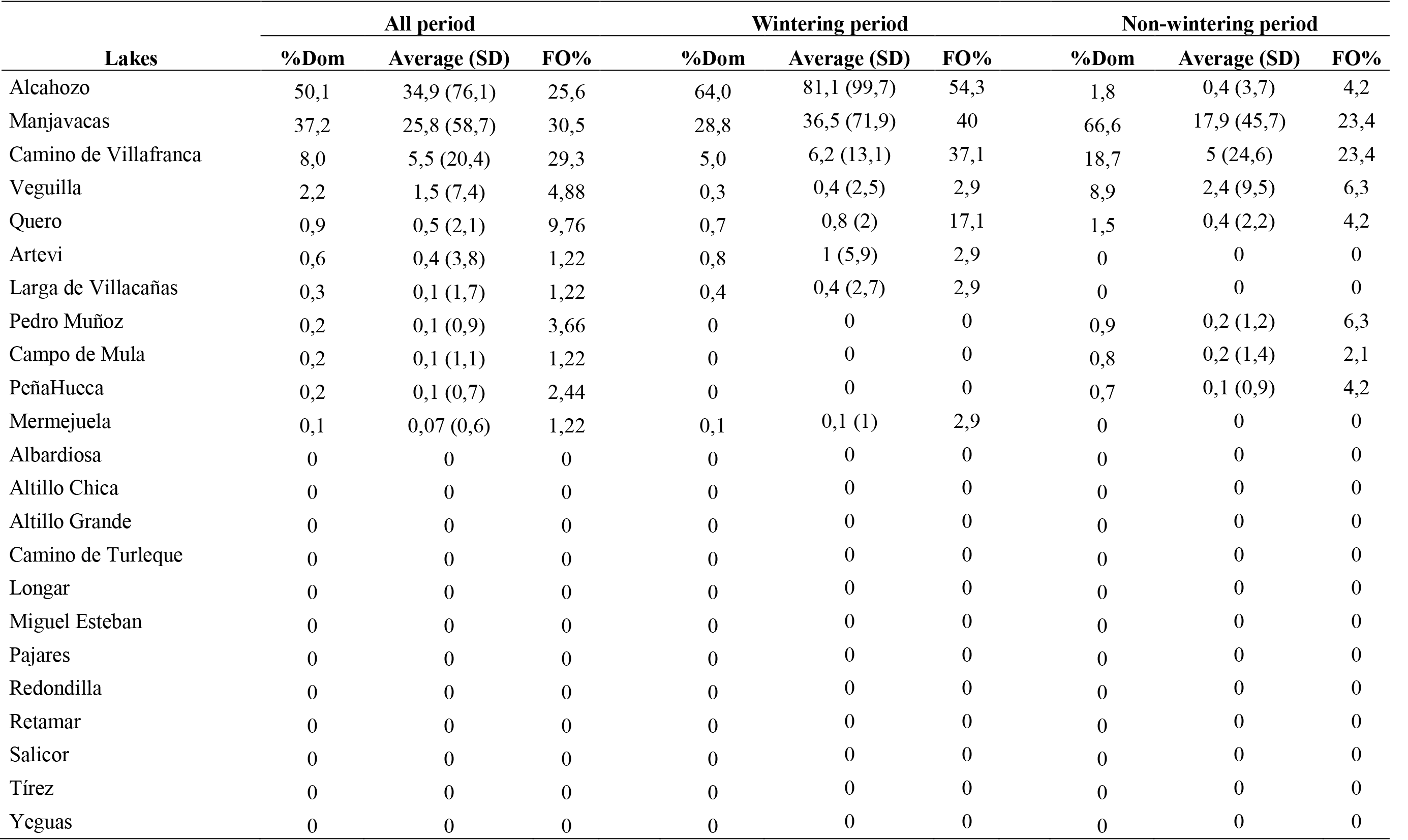
Lakes monitored between October 2010 and July 2017 and their respective dominance values (% Dom; distribution of accumulated records by period), average monthly abundance (average (SD)) and frequency of occurrence (FO%; number of months with presence of dunlin).

Temporal variations of abundance in two lakes concentrated most of the records, Alcahozo and Manjavacas, presented a low correlation (r = 0.3). In five of the seven winter seasons, abundance values reached the first peaks or occurred exclusively at Lake Alcahozo. Subsequently, abundance reductions in this lake were followed by an increase of individuals in Lake Manjavacas (Fig. 2).

**Fig. 2.**
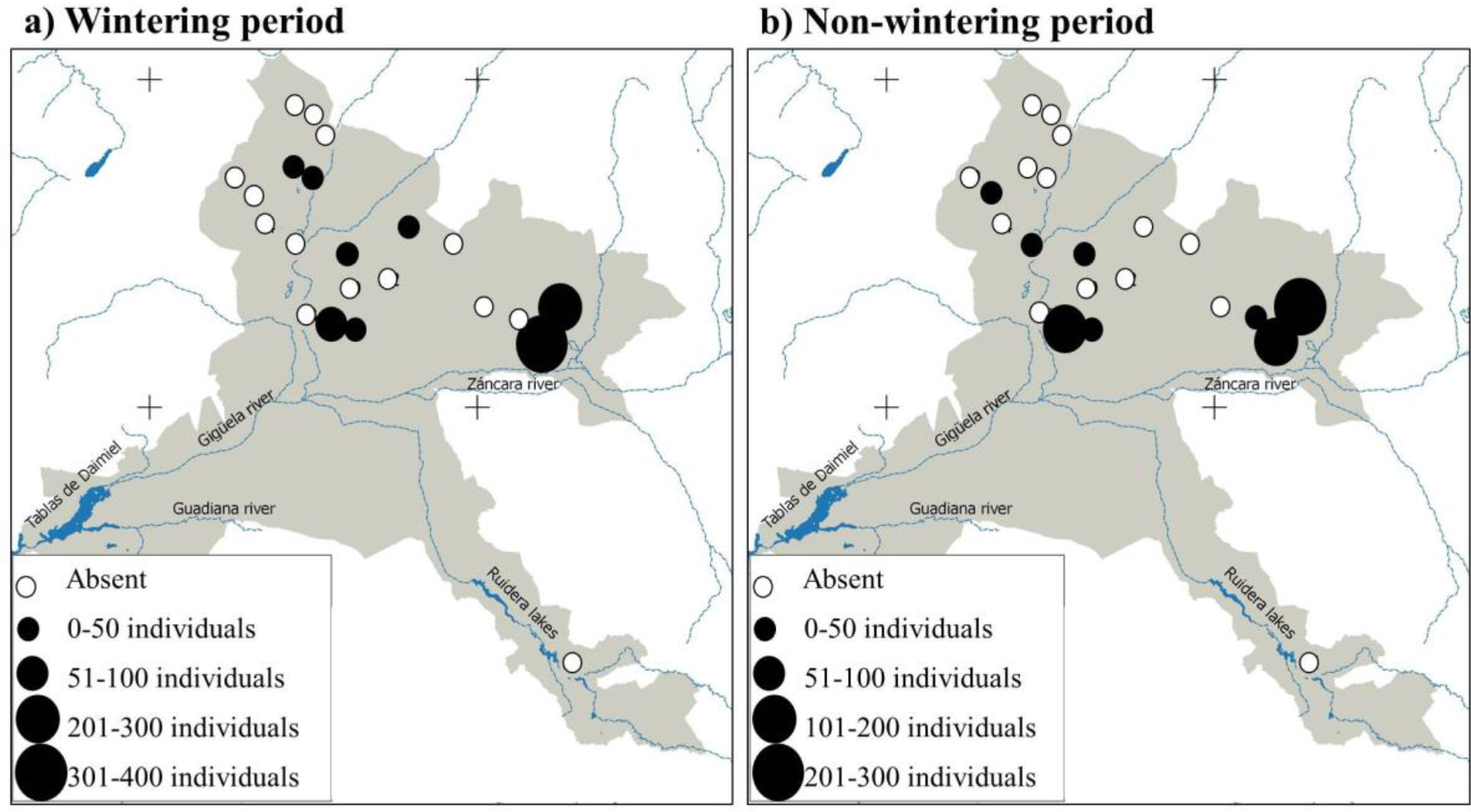
Distribution of the highest values of dunlin abundance observed in a single month for the wintering (a) and non-wintering (b) periods.

**Fig. 3.**
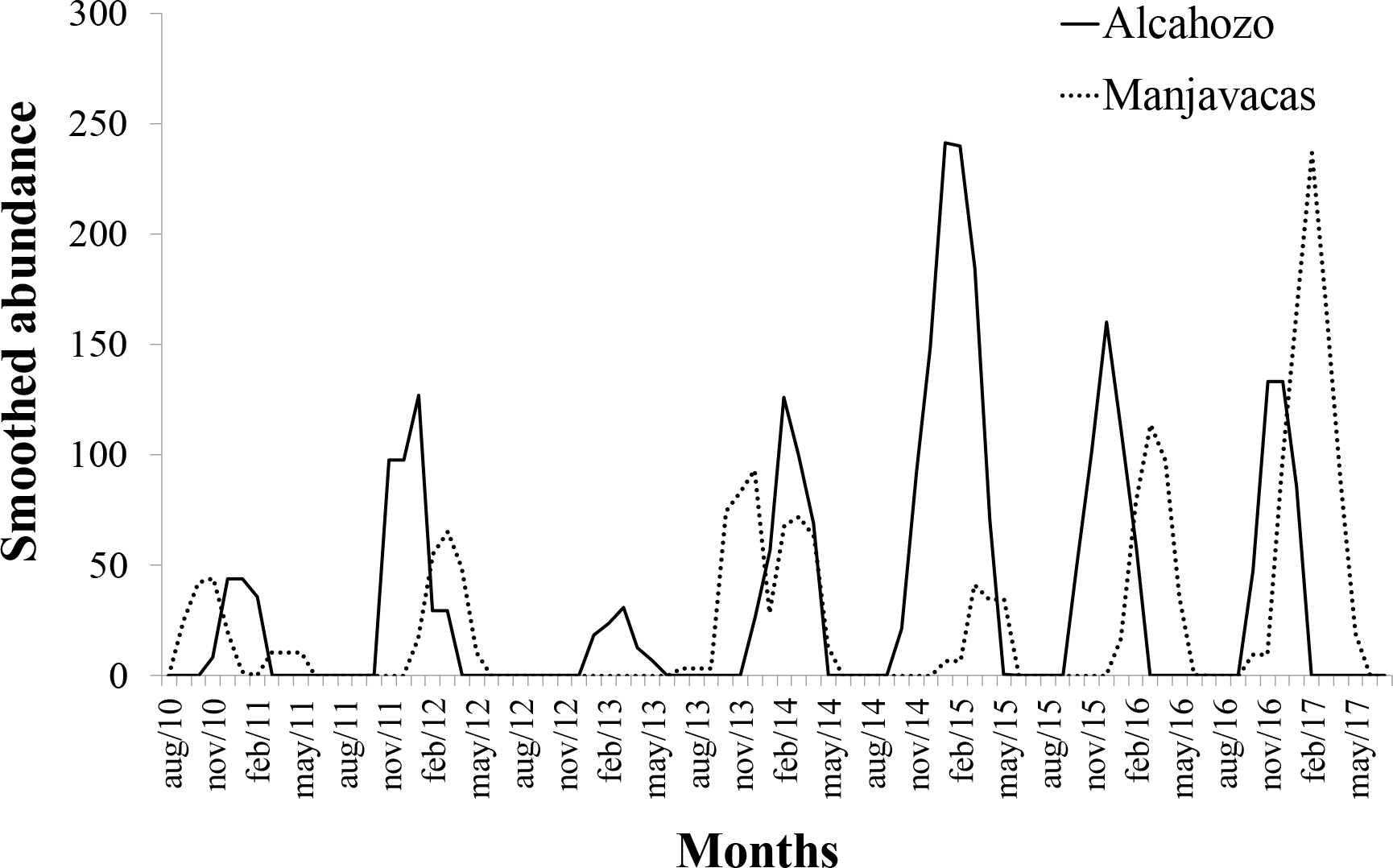
Temporal variation (moving averages) of the abundance of dunlin in two lakes of the Manjavacas Lagoon Complex.

## 4. Discussion

According to the atlas of birds in winter in Spain (Seo/BirdLife International 2012), which shows a retrospective of the annual abundance values of the migratory species, our data indicate that approximately 3% of the wintering population of dunlin is in our study areas. The values of FO and monthly average abundance for each pond per period indicate that few ponds concentrate the vast majority of individuals during the wintering and non-wintering periods.

Alcahozo and Manjavacas lakes, were the most important sites during the wintering period, accounting for almost 90% of the accumulated records. The lakes of this complex are marked by different hydrological dynamics (Gonçalves et al. 2016), with significant effects on quality of its waters and, consequently, on the structure of invertebrate communities. The only complex included in our study area that resembles these lakes is the Alcazar de San Juan Complex, which integrates the lakes Camino de Villafranca, Yeguas and Veguilla. However, this complex does not have a composition of invertebrates similar to the Manjavacas’ lakes (unpublished data from Ecolake Project, University of Valencia, Spain). Specifically, the set of invertebrate species of the lagoon complex of Manjavacas is marked by the abundance of copepods and anostracans (Pons et al. 2018). This is particularly important since the combination of anostracans and microcrustaceans has been referred to as key factors in the selection of foraging areas during the migratory cycle of waders (Horváth et al. 2013), which could explain such preference for the lakes Alcahozo and Manjavacas during the wintering, and, at least in part, for non-wintering period.

The temporal variations of abundance observed for Alcahozo and Manjavacas lakes indicate local movements of individuals within the Manjavacas complex as winter progresses. The first groups of wintering individuals tend to be observed first in the Alcahozo lake and, later, in the Manjavacas lake. This pattern may be associated with the aquatic invertebrate life cycle of both wetlands. Specifically, the main species of anostracan–*Branchinectella media*– is present only in Alcahozo lake. This species, besides being an indicator of environmental quality and being absent in polluted wetlands (Alonso 1996), such as the Manjavacas lake, is one of the first invertebrate species to appear at the beginning of the hydrological cycle (Jocque et al. 2010). Anostracans also present a fast growth and a high biomass when compared to the other invertebrate species occurring in Manjavacas lake (cladocers, copepods, and ostracods) (Alonso 1996). In this sense, the selection of the Alcahozo lake in early winter could indicate a response to the nutritional quality of prey, at least until the end of the life cycle of *B. media*, usually at the end of winter (Pons et al., 2017), when Dunlins begin to occupy the Manjavacas lake. This process could also explain the importance of the Manjavacas lake during the non-wintering period, when this lake remains part of time with water and presents great abundance of invertebrates (Boronat et al. 2001).

The recognition of spatiotemporal patterns of wintering populations helps to understand the factors that influence their habitat needs, as well as the recognition of the conservation status of their ecosystems (Lunardi et al. 2012). Our data help to understanding some of factors that influence of the distribution of dunlin during the wintering and non-wintering period in the “La Mancha Húmeda” Biosphere Reserve. Notably, we highlight that the main wintering area of the species – Manjavacas Lagoon Complex – is under strong anthropic pressure, because there is failures in the treatment of its wastewater, as well as under constant changes as a consequence of the surrounding agricultural activities. Finally, our data suggest that the lagoons belonging to this complex constitute of faithful wintering sites of the species and that conservation and improvement measures should be taken to keep viable the presence of these populations.

## References

Alonso, M. 1996. Crustacea, Branchiopoda. In: Fauna Ibérica, vol. 7. M. A. Ramos (ed.). MuseoNacional de CienciasNaturales, CSIC. Madrid. Spain.

Andres, B. A., Smith, P. A., Morrison, R. G., Gratto-Trevor, C. L., Brown, S. C., & Friis, C. A. 2012. Population estimates of North American shorebirds, 2012. Wader Study Group Bull, 119(3), 178–194.

BirdLife International. 2018 IUCN Red List for birds. Downloaded from http://www.birdlife.org on 11/09/2018.

Boronat, L., Miracle, M. R., & Armengol, X. 2001.Cladoceran assemblages in a mineralization gradient. Hydrobiologia, 442(1), 75–88.

Collins, S. L., & Glenn, S. M. 1991.Importance of spatial and temporal dynamics in species regional abundance and distribution. Ecology, 72(2), 654–664.

Crespo, J. G. C., García, M. A. R., & Bravo, A. L. 2011. Reserva de la Biosfera de la Mancha Húmeda: retos y oportunidades de futuro. Toledo: Dirección General de Áreas Protegidas y biodiversidad, Junta de Comunidades de Castilla La Mancha (in spanish).

Gonçalves, M. S. S., Gil-Delgado, J. A., Gosálvez, R. U., López-Iborra, G. M., Ponz, A., % Velasco, A. 2016. Spatial synchrony of wader populations in inland lakes of the Iberian Peninsula. Ecological Research, 31: 947–956.

Gonçalves, M. S., Gil-Delgado, J. A., Gosalvez, R. U., López-Iborra, G. M., Ponz, A., & Velasco, A. (in press). Seasonal differences in drivers of species richness of waders in inland wetlands of the “La Mancha Húmeda” Biosphere Reserve. Aquatic Conservation: Marine and Freshwater Ecosystems. doi:10.1002/aqc.2968.

Gosálvez, R. U., Gil-Delgado, J. A., Vives-Ferrándiz, C., Sánchez, G., & Florín, M. 2012.Seguimiento de aves acuáticasamenazadas em lagunas de la Reserva de la Biosfera de La Mancha Húmeda (Espanha Central). Polígonos, Revista de Geografía, 22, 89–122 (in spanish).

Green, A. J, & Elmberg, J. 2013. Ecosystem services provided by waterbirds.Biological Reviews, 89, 105–122.

Guadagnin, D. L, & Maltchik, L. 2007. Habitat and landscape factors associated with neotropicalwaterbird occurrence and richness in wetland fragments. Biodiversity and Conservation, 16, 1231–1244.

Guadagnin, D. L., Maltchik, L., & Fonseca, C. R. 2009. Species–area relationship of Neotropicalwaterbird assemblages in remnant wetlands: looking at the mechanisms. Diversity and Distribution, 15, 319–327.

Hammer,Ø., Harper D. A. T., Ryan P. D. 2001. PAST: Palaeontological Statistics software package for education and analysis, ver. 3.04. Palaeontologia Electronica 4:9.

Horváth, Z., Vad, C. F., Vörös, L., & Boros, E. 2013. The keystone role of anostracans and copepods in European soda pans during the spring migration of waterbirds. Freshwater Biology, 58(2), 430–440.

Jocque, M., Vanschoenwinkel, B. & Brendonck, L. 2010.Anostracanmonopolisation of early successional phases in temporary waters?. Fundamental and Applied Limnology/ArchivfürHydrobiologie, 176(2), 127–132.

Kingsford, R. T., Basset, A., & Jackson, L. 2016. Wetlands: conservation’s poor cousins. Aquatic Conservation: Marine and Freshwater Ecosystems, 26, 892–916

Lunardi, V. O., Macedo, R. H., Granadeiro, J. P., & Palmeirim, J. M. 2012. Migratory flows and foraging habitat selection by shorebirds along the northeastern coast of Brazil: the case of Baía de Todosos Santos. Estuarine, Coastal and ShelfScience, 96, 179–187.

Martinez-Santos P, De Stefano L, Llamas MR, Martínez-Alfaro PE (2008) Wetland restoration in the Mancha Occidental aquifer, Spain: a critical perspective on water, agricultural, and environmental policies. Restor Ecol 16:511–521 doi:10.1111/j.1526-100X.2008.00410.x

Ministerio de Agricultura, Alimentación y Medio Ambiente. 2014. Resolución de 17 de noviembre de 2014, de Parques Nacionales, por la que se publica la aprobación por la UNESCO de la ampliación de la Reserva de la Biosfera de Montseny, Cataluña, y la Reserva de la Biosfera de La Mancha Húmeda, en Castilla-La Mancha.

Morrison, R. I. G.; McCaffery, B. J.; Gill, R. E.; Skagen, S. K.; Jones, S. L.; Page, G. W.; Gratto-Trevor, C. L.; Andres, B. A. 2006. Population estimates of North American shorebirds, 2006. Wader Study Group Bull. 111: XX-XX.

Pons, P.; Gonçalves, M. S. S.; Ortells, R.; Gil-Delgado, J. A. 2017. Spatio-temporal population dynamics of Branchinectella media (Crustacea, Branchiopoda) from three saline ponds of the Iberian Peninsula.In 7th European Pond Conservation Network Workshop + LIFE Charcos Seminar.Algarve, Portugal. May 1- May 4. Abstract PST11.

Pons, P., Gonçalves, M. S. S., Gil-Delgado, J. A., & Ortells, R. 2018.Spatial distribution of Branchinectellamedia (Crustacea, Branchiopoda) in a saline pond from “La Mancha Húmeda”: a case of habitat selection?. Limnetica, 37: 69–83

Ruiz-Pulpón, A. R. 2013. El viñedo en espaldera: nueva realidad en los paisajes vitivinícolas de Castilla-La Mancha. Boletin de la Associación de Geógrafos Españoles, vol: 249–270.

Ruiz-Pulpón, A. R. 2015. Dinámicas de mercado y transformación de los paisajes vitivinícolas de Castilla-La Mancha. In: Análisis espacial y representación geográfica: innovación y aplicación: 2141–2150 [J. De la Riva, P. Ibarra, R. Montorio, M. Rodrigues, Eds.]. Universidad de Zaragoza – AGE, Zaragoza.

SEO/BirdLifeInternational. 2012. Atlas de las aves en invierno en España 2007-2010. Ministerio de Agricultura, Alimentación y Medio Ambiente - SEO/BirdLife. Madrid, pp 234–235 (in Spanish)

